# Evaluation of podocyte Rac-1 induced kidney disease by modulation of TRPC5

**DOI:** 10.1101/2021.05.25.445694

**Authors:** Onur K. Polat, Elena Isaeva, Ke Zhu, Manuel Noben, Yashwanth Sudhini, Beata Samelko, Varsha S. Kumar, Changli Wei, Mehmet M. Altintas, Stuart E. Dryer, Sanja Sever, Alexander Staruschenko, Jochen Reiser

## Abstract

**Background:** Transient receptor potential channel 5 (TRPC5) is a non-selective cationic ion channel expressed in brain, kidney and other organs where its activation underlies podocyte injury in chronic kidney diseases. Specifically, it has been suggested that a podocyte TRPC5 plasma membrane relocation and channel activation following injury results from activation of Rac-1, propagating podocyte dysfunction and proteinuria. However, previous TRPC5 transgenic mouse studies had questioned a pathogenic role for TRPC5 in podocytes. This investigation was designed to specifically evaluate podocyte Rac-1 activation in the context of functional TRPC5 or a TRPC5 pore mutant to assess effects on proteinuria.

**Materials and Methods:** We employed single cell patch-clamp studies of cultured podocytes and studied proteinuria in transgenic mouse models to characterize the effects of TRPC5 following podocyte Rac-1 activation.

**Results:** Inhibition of TRPC5 by small molecules reportedly ameliorated proteinuria in murine models of proteinuric kidney diseases. In order to directly examine TRPC5 function following Rac-1-induced podocyte injury, we analyzed TRPC5 inhibition in podocyte specific Rac-1 (active) transgenic mice. In addition, we generated a double-transgenic mouse constitutively overexpressing either TRPC5 (TRPC5^WT^) or a TRPC5 dominant-negative pore mutant (TRPC5^DN^) in concert with podocyte specific and inducible activation of active Rac-1 (Rac-1^Dtg^). In electrophysiological experiments, active TRPC5 was detected in primary podocytes overexpressing TRPC5 but not in podocytes with endogenous TRPC5 expression, nor with Rac-1 overexpressing podocytes. TRPC5 inhibition did not change proteinuria in mice with active podocyte Rac-1, nor did an increase or loss of TRPC5 activity affected podocyte injury in Rac-1^Dtg^ animals. Administration of TRPC5 inhibitors, ML204 and AC1903, did not alleviate podocyte Rac-1 induced proteinuria.

**Conclusion:** TRPC5 inhibition did not modify podocyte Rac-1 induced proteinuria in mice.

**Significance Statement:** TRPC5 is a calcium conducting ion channel involved in a plethora of biological functions in the brain, kidney and other organs. In proteinuric kidney diseases, others proposed a model that links activation of small GTPase Rac-1 in podocytes to activation of TRPC5 channels propagating cellular injury and eventually leading to progressive kidney disease. To test this hypothesis, we have developed a novel transgenic mouse model that employs podocyte Rac-1 activation in the presence or absence of a functional TRPC5 channel. Our data shows that transgenic mice with activated Rac-1 in podocytes did not enhance endogenous TRPC5 expression or its activity. Furthermore, TRPC5 blockade or activation did not modify Rac-1 induced proteinuria in mice.

## Introduction

Transient receptor potential canonical channels (TRPCs) are non-specific cationic ion channels expressed in various kidney segments, which play a role in signal transduction, especially in G-protein signaling cascades. The first TRPC identified within the podocytes of the kidneys was TRPC6, where it was shown that “gain of function” mutations within *TRPC6* led to a variant of chronic kidney disease (CKD) termed as focal segmental glomerulosclerosis (FSGS)^1,2,3^. Subsequent experiments have suggested that TRPC6 may also play a role in driving acquired forms of CKD. While kidney disease associated with loss-of-function *TRPC6* mutations has also been reported^4^, it is generally thought that chronic dys-regulated Ca^2+^-influx from TRPC6 activity leads to podocyte injury underlying progressive CKD and familial FSGS^5^.

TRPC5 is a receptor-stimulated ion channel that can also be detected in the kidney. It has been suggested that TRPC5-mediated Ca^2+^ influx induces Rac-1 activation leading to cytoskeletal dysregulation, which in glomerular podocytes underlies foot processes effacement and proteinuria. The actin cytoskeleton in podocytes is regulated by both RhoA and Rac-GTPases and their associated Ca^2+^ channels such as TRPC6 and TRPC5^6^. In contrast to TRPC5 mediated currents, TRPC6-mediated Ca^2+^ influx was linked to RhoA activation and decreased cell motility. It has been suggested that TRPC5 and TRPC6 work in tandem to manage the global actin cytoskeleton dynamics in podocytes^6,7,8^.

It is postulated that injurious events to podocytes activates Rac-1, but also induces translocation of the TRPC5 channels from the storage vesicles in the cytosol to the plasma membrane (Fig. 1A)^9^. TRPC5-mediated influx of Ca^2+^ subsequently leads to further Rac-1 activation which in turn locks TRPC5 on the plasma membrane^10,11^ This model predicts that disrupting the connection between TRPC5 and Rac-1 would eliminate the Ca^2+^ influx after Rac-1 activation, thus ending the positive feedback loop and protecting podocytes from Rac-1 induced proteinuria (Fig. 1A).

**Figure:**
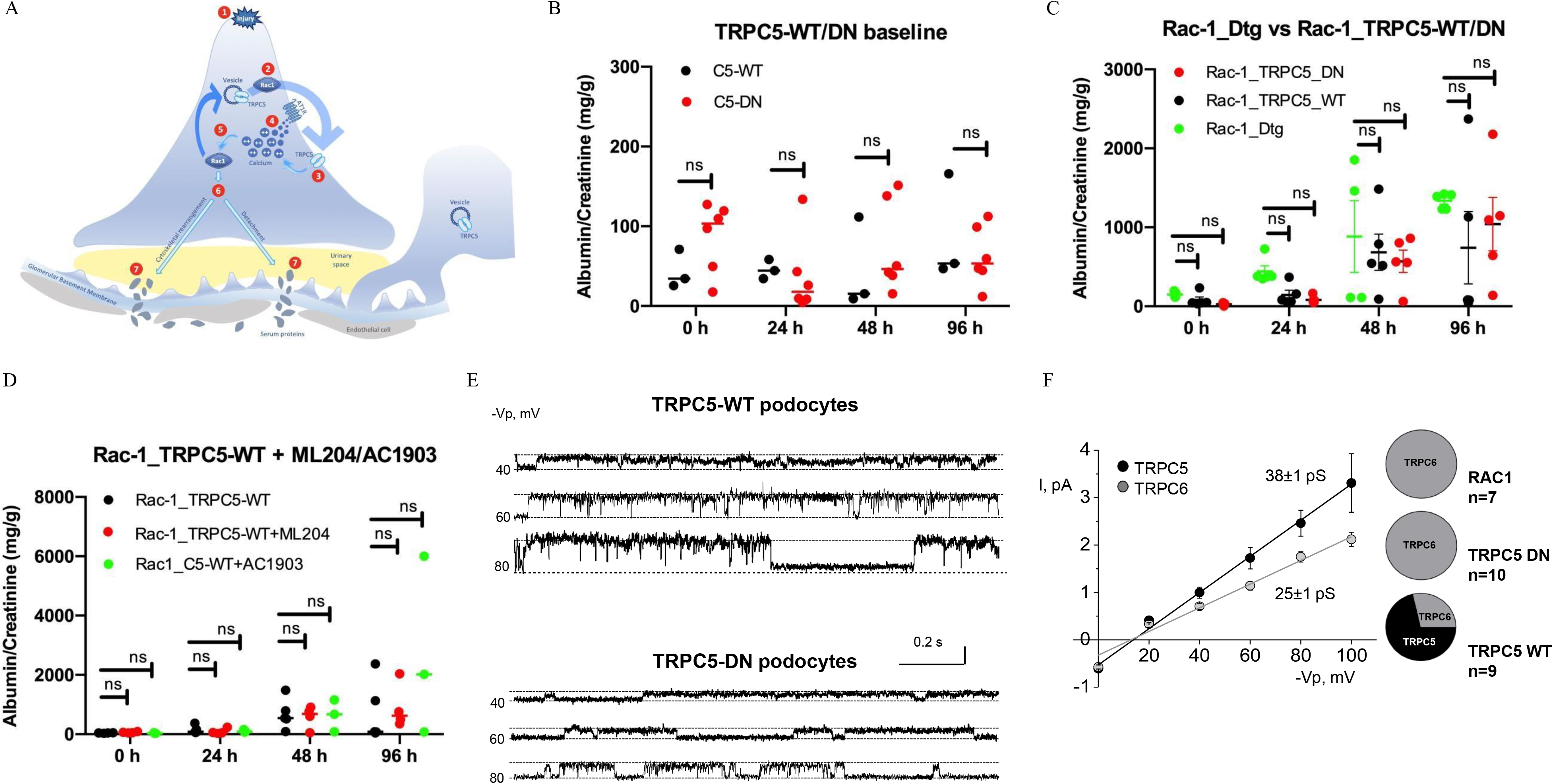
TRPC5 activity does not influence Rac1-induced proteinuria. **(A)** model adapted from Chung and Shaw^11^. The model depicts the endless feedback loop betweenTRPC5 and Rac-1. The sequence of events (1)-Injury due to external or internal factors, (2)-Rac-1 is upregulated, (3)-TRPC5 is brought to the surface due to the presence of Rac-1, (4)-Ca^2+^ influx occurs, (5)-Ca^2+^ influx leads to Rac-1 activation, (6)-Activated Rac-1 signals for mediator molecules, (7)-Prolonged exposure to signaling molecules lead to damage to the podocytes. **(B)** Changes in albumin (mg) to creatinine (g) ratio (ACR) over time following of TRPC5^WT^ and TRPC5^DN^ with regular chow. The black dots indicate TRPC5^WT^ mice whereas red dots show the data for TRPC5^DN^ mice. Error bars indicate SEM. No significance was detected betweengroups (p>0.05; NS for all cases measured by unpaired-students t-test following satisfactory normality tests). **(C)** Albumin (mg) to creatinine (g) ratio (ACR) was shown on the y-axis with the day-by-day progression of the proteinuria across mice groups shown on the x-axis. The green plot indicates Rac-1^Dtg^ mice, the black color plot indicates Rac-1^Dtg^─TRPC5^WT^ mice, and the red color shows Rac-1^Dtg^─TRPC5^DN^ mice. Error bars indicate SEM. No significance was detected between groups (p>0.05 for all cases, measured by *one-way* ANOVA followed by Bonferroni post-hoc correction). **(D)** ACR measurements of mice injected with PBS (black bars), ML204 (red bars), and AC1903 (green bars). Doses of ML204 and AC1903 were 0.3 and 0.75 mg/kg, respectively. Error bars are shown as SEM. No significance was detected between groups (p>0.05 for all cases, measured by *one-way* ANOVA followed by Bonferroni post-hoc correction). **(E)**Representative traces of ionic currents demonstrating activity from TRPC5^WT^ and TRPC5^DN^ podocytes are shown. Rac-1^Dtg^ representative trace is identical to the trace of TRPC5^DN^, and is not shown separately, however is included in the final analysis in Fig. 1F. All animals were genotyped with appropriate protocols before experimentation. **(F)** Current-voltage (I/V) relationships for identified channels. Two types of channels were detected in cells from mice overexpressing TRPC5, with unitary slope conductances of 38±1 pS, and 25±1 pS. However, only the 25 pS channels were detected in mice overexpressing TRPC5^DN^ or Rac-1^Dtg^. The pie chart depicts the relative abundance of the two types of unitary currents in the patches.

## Materials and Methods

### Mice

All animal experiments and protocols were approved by the Rush University Institutional Animal Care and Use Committee. Food and water were available *ad libitum*. TRPC5-wild-type (TRPC5^WT^) and TRPC5-dominant negative (TRPC5^DN^) over-expressing mice were generated as described earlier^12^. Podocyte-inducible Rac-1 mice (Rac-1^Dtg^) were a gift from Dr. Andrey Shaw (Washington University in St. Louis). Rac-1^Dtg^ and TRPC5^WT/DN^ mice were crossed to obtain the Rac-1^Dtg^─TRPC5^WT/DN^ double transgenic mice. All mice genotypes were confirmed with genomic PCR.

### Proteinuria induction

Proteinuria was induced by activating Rac-1, with addition of DOX-laced chow (2000 μM) *ad libitum* for 4 days as described before^13^. For baseline controls TRPC5^WT/DN^ were fed normal chow.

### Urine collection

Urine were collected from each mouse within each group for 4 days, every 24 hours. Urine samples were stored at −80°C until further use.

### Albumin to creatinine ratio (ACR)

Urinary ACR was measured by Albumin Elisa kit (Bethyl Laboratories, Montgomery, TX) and Creatinine assay kit (Cayman Chemicals, Ann Arbor, MI). Collected urines were diluted 1:20 (for the creatinine assay) and 1:5000 (for the albumin assay) and assayed based on the manufacturer’s recommendations. Obtained albumin and creatinine values were converted to mg/mL and g/mL and ratioed. ACR with a value of more than 100 was considered proteinuric.

### Genomic PCR

Tails of each mice were cut and placed in the digestion buffer (MCG buffer, 10% Triton-X, **β**-mercaptoethanol and dH_2_O). After 3 min of boiling at 93°C, the tails were allowed to cool to room temperature, followed by the addition of Proteinase K. The digestion mixture was left to digest overnight at 60°C. Proteinase K was neutralized by heat induction (95°C for 5 min) and 1 μl crude DNA lysate was amplified with the *Taq-Blue* PCR kit according to the instructions from the manufacturer. The DNA was amplified with the following gene specific primers:

<u>TRPC5 WT/DN primer:</u> *5’-TGCAAACATCACATGCACAC -3’*
<u>Rac-1 primer:</u> *5’-GAAGCAGAAGCTTAGGAAGATGG-3’*

Resulting amplifications were subjected to 1% agarose-gel and were visualized using SYBR-safe DNA gel stain.

### ML204 and AC1903

DOX-dependent proteinuria was achieved as described above. TRPC5 inhibitors, 0.3 mg/kg ML204 (176 μM) or 0.75 mg/kg AC1903 (333 μM), were dissolved in dimethylsulfoxide (DMSO) and injected into mice 48 and 76 h after DOX-chow addition to establish the effect of ML204 and AC1903 (both from Sigma-Aldrich, St. Louis, MO) on ACR, respectively.

### Glomeruli isolation and primary podocyte generation

Mice were euthanized by CO_2_ overdose following cervical dislocation. Immediately after, the kidneys were collected, rinsed of blood and decapsulated. The cortex was minced with a steel blade on a petri dish filled with ice-cold sterile PBS. The resultant minced slurry was passed through three stack metal sieves with 100, 70 and 53 μm meshes. The slurry was washed thoroughly with ice-cold PBS and the material that remained in the 53 μm mesh was collected. The collected material was then centrifuged at 1200 rpm for 8 min. The pellet was resuspended in appropriatevolume of primary podocyte medium (RPMI 1640 supplemented with 10% FBS, 100 U/mL penicillin, 100 μg/mL streptomycin). Glomeruli were left undisturbed with intermittent medium changes for 14 days to which they differentiated to primary podocytes.

### Electrophysiology

Isolated podocytes were cultured for 14 days and subjected to single-channel patch-clamping as described previously^14^.

### Statistics

Statistical analysis was performed using Prism 9.0 software (GraphPad, La Jolla, CA). *Students t-test* was employed when comparing two groups to each other, after satisfying normality distribution conditions. All other significance tests were *one-way* ANOVA with Bonferroni *post-hoc* correction. *p*>0.05 (N.S.), *p*<0.05 (*), *p*<0.01(**), *p*<0.001 (***).

## Results and Discussion

To directly test the model of a Rac-1 induced TRPC5 feed forward loop originally suggested by Zhou et al. (Fig. 1A), we generated double transgenic mice that over-express either TRPC5^WT^ or TRPC5^DN^ mice (TRPC5 containing a dominant-negative mutation in the pore domain of the channel), together with podocyte specific and inducible activation of active Rac-1 (Rac-1^Dtg^) under the doxycycline (DOX) promoter. These mouse models clarified two important points currently under scientific debate. Firstly, podocyte Rac-1 activation does not lead to an enhanced activity of TRPC5 channel and secondly, having more or less TRPC5 activity does not influence proteinuria induced by Rac-1 activation in podocytes.

Baseline ACR values of TRPC5^WT^ and TRPC5^DN^ were initially evaluated to rule out stress or handling-related changes to the urine protein level. Mice were fed normal chow for 4 days and urine was collected at 0, 24, 48, and 96 hours. The urine was subjected to albumin:creatinine ratio (ACR) analysis, which is expressed in mg/g, and an ACR value greater than 100 mg/g is considered to indicate proteinuria. While some mice exhibited higher than expected baseline ACR values, proteinuria did overall not progress within the observed 4 days suggesting absence spontaneous proteinuria (Fig. 1B).

We next examined whether our novel transgenic mouse models (Rac-1^Dtg^, Rac-1^tg^─TRPC5^WT^, and Rac-1^Dtg^─TRPC5^DN^) developed proteinuria. When these mice were fed DOX-laced chow for 4 days, as described above, Rac-1^Dtg^,Rac-1^Dtg^─TRPC5^WT^ and Rac-1^Dtg^─TRPC5^DN^ mice developed significant proteinuria at 48 h, which reached a peak at 96 h (Fig. 1C). There were no significant differences in urine albumin excretion between Rac-1^Dtg^, Rac-1^Dtg^─TRPC5^WT^ and Rac-1^Dtg^─TRPC5^DN^ mice (Fig. 1C). These data strongly suggested that neither increases nor decreases in functional activity of podocyte TRPC5 affect glomerular protein filtration associated with sustained activation of Rac-1. These results are also consistent with our previous study in which englerin A, an activator of TRPC4 and TRPC5, did not increase proteinuria^12^.

To further assess the role of TRPC5 in Rac-1 induced proteinuria, we examined the effects of ML204 and AC1903, two structurally different small molecules that can inhibit TRPC5^10,15^. Following baseline measurements (at t=0 h), Rac-1^Dtg^─TRPC5^WT^ mice were placed on DOX-laced chow. Proteinuria was detected as early as 24 h (ACR>100 mg/g). At 48 and 72 hours, the mice were injected by *i.p.* route with either a vehicle (e.g., PBS) or 0.3 mg/kg ML204 or 0.75 mg/kg AC1903. We did not observe a statistically significant difference between mice injected with PBS or mice treated with either ML204 or AC1903. In fact, Rac-1^Dtg^─TRPC5^WT^ mice treated with AC1903 showed greater urine albumin excretion after 96 h, where the Rac-1 induced proteinuria appears to be the most severe (Fig. 1D).

To confirm the functional expression and activity of TRPC5, we isolated primary podocytes and evaluated TRPC channel activity in the cell-attached mode of patch-clamp configuration. Primary podocytes derived from two TRPC5^WT^, TRPC5^DN^, and Rac-1^Dtg^ animals exhibited two distinct types of channels based on their unitary conductance. The higher conducting current (~38±1 pS; n=6) was only detected in podocytes from mice over-expressing TRPC5^WT^, whereas the second type of channel (~25±1 pS; n=18) was detected in all three lines of transgenic mice (Figs 1E and 1F). Based on previous studies demonstrating that the unitary conductance of TRPC5 is higher than that of TRPC6^16,17,18,19^, and our studies utilizing TRPC6 knockout mice^14^, we can conclude that the recorded channels with 38 and 25 pS conductance represent activity of TRPC5 and TRPC6, respectively (Figs 1E and 1F). TRPC5 currents were only detected in cultures of TRPC5^WT^ podocytes but not in TRPC5^DN^ (Fig. 1E) or Rac-1^Dtg^ podocytes (data not shown). Currents detected from Rac-1^Dtg^ and TRPC5^DN^ resembled TRPC6 currents as summarized in Fig. 1F). These data show that active TRPC5 channels on the podocyte cell surface are only present in TRPC5-overexpressing mice and that over-expression of Rac-1^Dtg^ was not sufficient to lead to activation of endogenous TRPC5 and cause the positive feed-back loop between Rac-1 and TRPC5 as the models suggest.

In summary, these data suggest that changes in TRPC5 do not affect the glomerular function and do not appear to play a role in kidney pathology caused by sustained activation of Rac-1. It should be noted that sustained activation of Rac-1 would be expected to produce various other effects in cells, for example, activation of NADPH-oxidase-2 (NOX2), generation of reactive oxygen species, and activation of TRPC6^5^.

## Funding

This work was supported by Department of Veteran Affairs Grant I01 BX004024 and the National Institutes of Health Grants R35 HL135749.

## Author contributions

O.K.P., E.I., K.Z., M.N., S.Y., B.S. performed experiments. V.S.K., C.W., M.M.A. performed data analysis, S.E.D., S.S., A.S., and J.R. critiqued, commented, and wrote the paper. All authors commented, revised and approved the manuscript

## Competing Interests

S.S. and J.R. are co-founders and shareholders of Walden Biosciences, a biotechnology company that develops novel kidney-protective therapies.

